# Divergent evolution of mutation rates and biases in the long-term evolution experiment with *Escherichia coli*

**DOI:** 10.1101/2020.06.02.130906

**Authors:** Rohan Maddamsetti, Nkrumah A. Grant

## Abstract

All organisms encode enzymes that replicate, maintain, pack, recombine, and repair their genetic material. For this reason, mutation rates and biases also evolve by mutation, variation, and natural selection. By examining metagenomic time series of the Lenski long-term evolution experiment (LTEE) with *Escherichia coli* (Good, et al. 2017), we find that local mutation rate variation has evolved during the LTEE. Each LTEE population has evolved idiosyncratic differences in their rates of point mutations, indels, and mobile element insertions, due to the fixation of various hypermutator and antimutator alleles. One LTEE population, called Ara+3, shows a strong, symmetric wave pattern in its density of point mutations, radiating from the origin of replication. This pattern is largely missing from the other LTEE populations, most of which evolved missense, indel, or structural mutations in *topA*, *fis*, and *dusB*— loci that all affect DNA topology. The distribution of mutations in those genes over time suggests epistasis and historical contingency in the evolution of DNA topology, which may have in turn affected local mutation rates. Overall, the replicate populations of the LTEE have largely diverged in their mutation rates and biases, even though they have adapted to identical abiotic conditions.

## INTRODUCTION

Loci that modify DNA repair and recombination modify the evolutionary process. Therefore, one might ask whether natural selection adaptively tunes mutation and recombination rates. This idea— that second-order selection *adaptively* modifies the evolutionary process itself— is debated (Tenaillon, et al. 2001; Lynch, et al. 2016). Nonetheless, populations of *Escherichia coli*, engineered to have constitutive sexual recombination and elevated mutation rates, adapt faster than control populations in the laboratory (Cooper 2007; Peabody, et al. 2016; Peabody, et al. 2017).

In order to study second-order selection on mutation rates, one can use experimental evolution. By running experiments in which replicate populations evolve under controlled conditions, with different starting mutation rates, one can ask whether particular mutation rates are favored over others (Chao, et al. 1983; Loh, et al. 2010; Sprouffske, et al. 2018). Here, we use metagenomic time series data from the Lenski long-term evolution experiment with *Escherichia coli* (LTEE) to study how mutation rates evolve in real-time.

In the LTEE, 12 populations of *Escherichia coli,* descended from a common ancestral strain, have adapted for more than 73,000 generations to carbon-limited minimal media. The LTEE populations are strictly asexual. Some populations have evolved defects in DNA repair which vastly increase their mutation rate. Those hypermutator alleles likely went to fixation by linkage with highly beneficial mutations, rather than being beneficial *per se* (Sniegowski, et al. 1997; Tenaillon, et al. 2016).

Molecular evolution in the hypermutator populations of the LTEE is dominated by “genetic draft”, in which large numbers of nearly neutral passenger mutations hitchhike with a small number of beneficial driver mutations. This phenomenon has obscured the genomic signatures of adaptation in those populations (Tenaillon, et al. 2016; Couce, et al. 2017; Good, et al. 2017; Maddamsetti, et al. 2017). In this regime, also called “emergent neutrality” (Schiffels, et al. 2011), the evolutionary dynamics inferred from whole-population samples of the hypermutator populations (Good et al. 2017) provides good data on mutation rates and biases, even though natural selection drives the dynamics. Here, we examined LTEE metagenomics data (Good, et al. 2017) for the mutation biases that have been reported by several research groups, albeit under different experimental conditions and with different strains (Foster, et al. 2013; Paul, et al. 2013; Jee, et al. 2016; Niccum, et al. 2019).

## RESULTS

### Cumulative number of observed mutations in each population reveals dynamics caused by both hypermutator and antimutator alleles

We examined the number of observed mutations over time in each LTEE population (Figure 1, Supplementary Figures S1–S3). These results show that mutation rates have evolved idiosyncratically over the LTEE. Figure 1A shows the number of point mutations over time in each population. The rate of observed point mutations decreased in three of the six hypermutator populations (Ara−2, Ara+3, and Ara+6). The decrease in the rate of molecular evolution in these populations was previously ascribed to the evolution of antimutator alleles (Tenaillon, et al. 2016; Good, et al. 2017). While antimutator alleles of *mutY* compensating for defects in *mutT* have been reported in Ara–1 (Wielgoss, et al. 2013), the change in slope seen around 40,000 generations in Ara−1 is subtle compared to the changes in slope seen in Ara−2, Ara+3, and Ara+6. The antimutator allele in Ara−2 will be discussed shortly, while those in Ara+3 and Ara+6 remain unknown.

**Figure 1.**
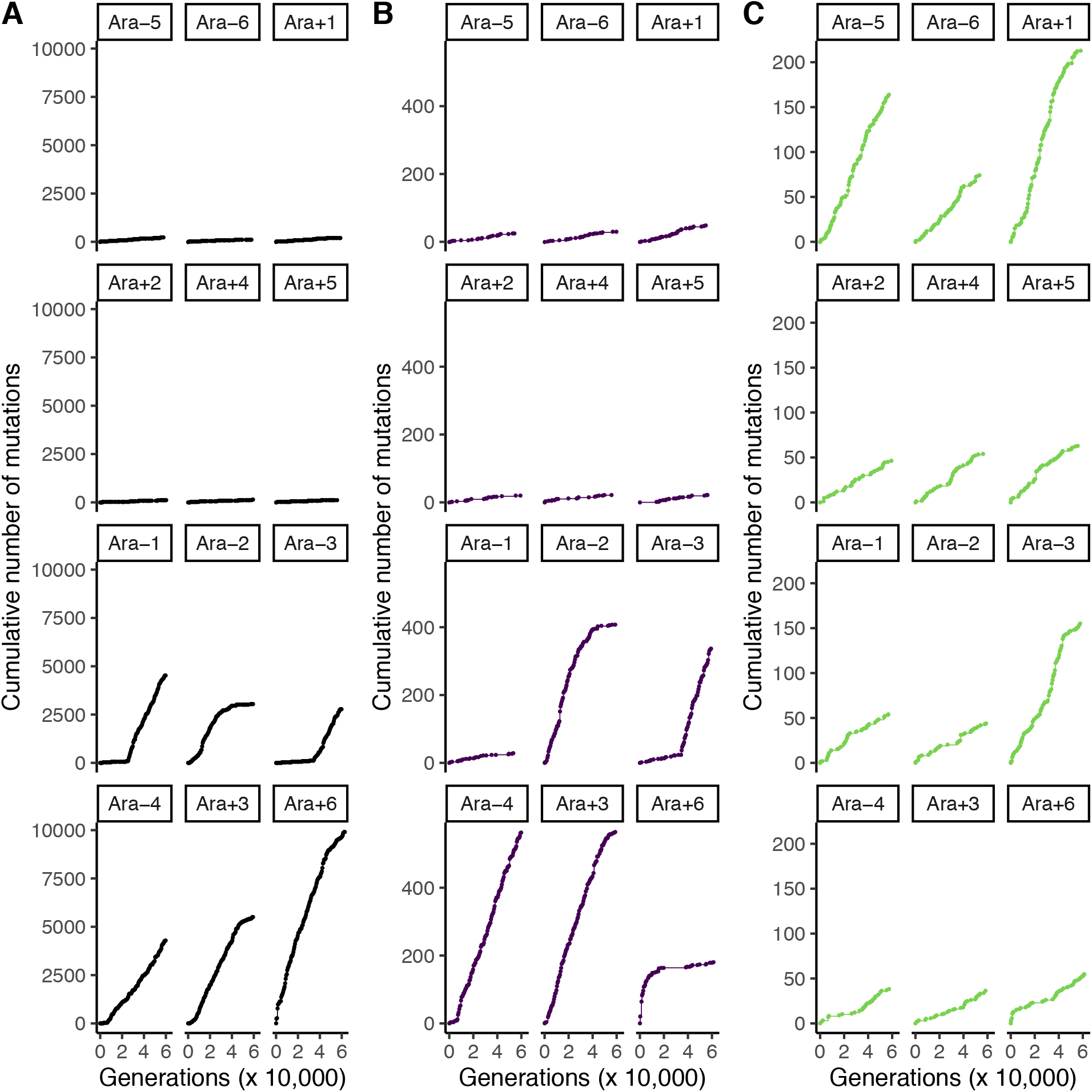
Divergent evolution of mutation rates in the LTEE. Each panel shows the cumulative number of observed mutations, subdivided by mutation class, over time in each LTEE population. The top six panels show the nonmutator LTEE populations, and the bottom six panels show the hypermutator LTEE populations. A) Point mutations are shown in black. B) Indel mutations are shown in purple. C) Structural mutations are shown in green.

In examining *mutT*, we noticed that two of the three cases of *mutT* alleles arising to high frequency in the LTEE occur on a *uvrA* background (Ara−2 and Ara+6), while the third, in Ara−1, occurs on a *uvrC* background (Figure 2). Might this indicate epistasis between *mutT* and *uvrABC*? It has been reported that *uvrA/mutT* and *uvrB*/*mutT* double mutants have a substantially lower mutation rate than a *mutT* mutant in the presence of hydrogen peroxide (Hori, et al. 2007). Thus, it is possible that the *mutT* alleles that successfully went to fixation in the LTEE evolved on an epistatic genetic background that reduces the intensity of their mutator phenotype.

**Figure 2.**
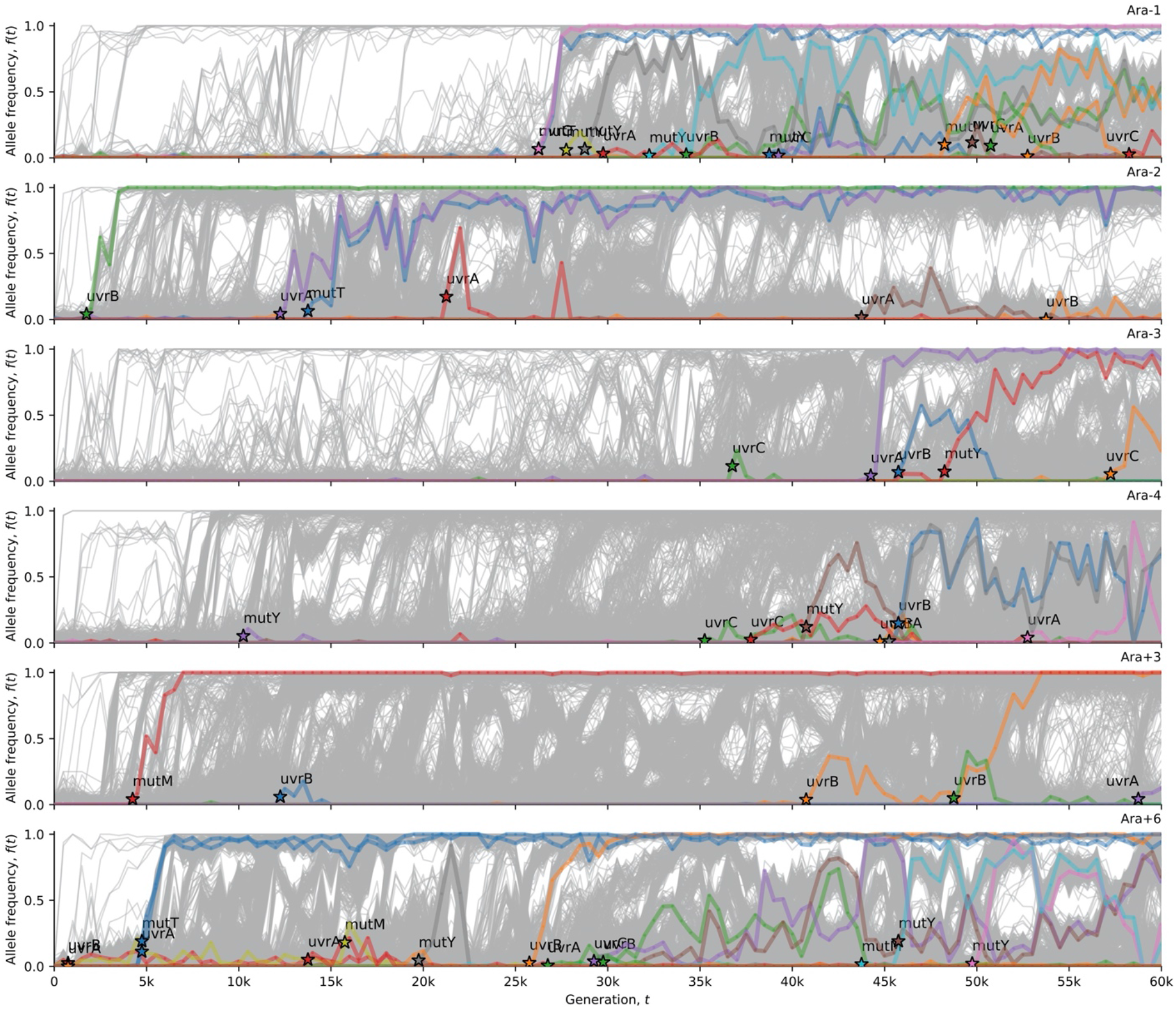
Oxidative damage repair alleles in hypermutator LTEE populations. This visualization uses computer code from Good et al. (2017). Stars indicate the time (and allele frequency) at which mutations are reliably estimated to appear in the time series. The allele frequency trajectories for all observed mutations in the hypermutator populations are shown in grey. The allele frequency trajectories of *de novo* mutations (excepting synonymous mutations) in oxidative damage repair genes (Supplementary File 1) are colored and labeled in each population.

Figure 1B shows the number of observed indel mutations over time in each population. 5 of the 6 point-mutation hypermutator populations also show an indel hypermutator phenotype. These 5 populations all evolved defects in mismatch repair (Figure 3). The exception is Ara−1, which evolved alleles of *uvrA* and *mutT* (Figure 2) that cause a high rate of point mutations, without a corresponding indel hypermutator phenotype.

**Figure 3.**
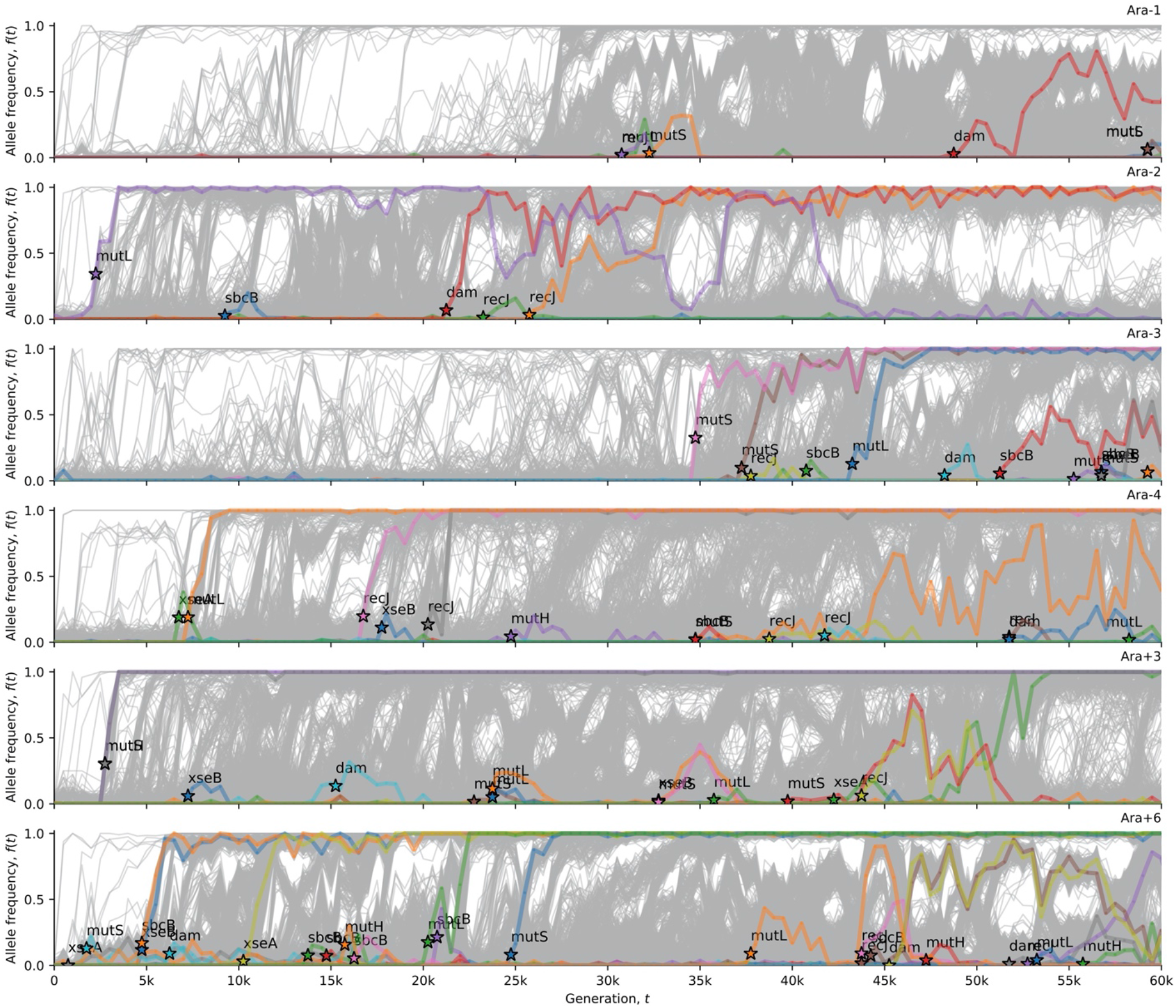
Mismatch repair alleles in the hypermutator LTEE populations. This visualization uses computer code from Good et al. (2017). Stars indicate the time (and allele frequency) at which mutations are reliably estimated to appear in the time series. The allele frequency trajectories for all observed mutations in the hypermutator populations are shown in grey. The allele frequency trajectories of *de novo* mutations (except synonymous mutations) in mismatch repair genes (Supplementary File 1) are colored and labeled in each population.

The hypermutator dynamics in Ara−2 are particularly striking. An antimutator allele eventually fixes, and reverts both the point and indel hypermutator phenotype back to wild-type or near wild-type levels (Figure 1A and 1B). The hypermutator phenotype is caused by phase variation of a (TGGCGC)_3_ repeat in *mutL*. Reversions to the triplet state reverse the hypermutator phenotype. The number of new point and indel mutations in Ara−2 (Supplementary Figures S1, S2) fluctuates with the allele frequency dynamics of this *mutL* repeat (Figure 3).

At first glance, Figure 1B seems to show that Ara+6 fixed a mutation reverting the indel hypermutator phenotype. However, the allele frequency dynamics reveal that a super-hypermutator clade evolved within the first 1000 generations. This super-hypermutator clade carried alleles of the mismatch repair genes *xseA* and *mutS* (Figure 3) as well as alleles of the nucleotide excision repair genes *uvrA* and *uvrB* (Figure 2), and persisted at low frequency until going extinct by 20,000 generations (Supplementary Figure S2). The majority clade in Ara+6 evolved mutations in *uvrA* and *mutT* at ~5000 generations (Figure 2) that caused a point mutation hypermutator phenotype without causing an indel hypermutator phenotype. The coexistence of clades with different hypermutator phenotypes, and the eventual extinction of the super-hypermutator clade, best explains the loss of the indel hypermutator phenotype from Ara+6.

Figure 1C shows the number of observed structural mutations over time. Most of these structural mutations are caused by the transposition of mobile genetic elements called insertion sequences (IS). Three of the canonical nonmutator populations (Ara−5, Ara−6, and Ara+1) show an IS hypermutator phenotype. The IS hypermutator phenotype in Ara+1 has been reported previously (Papadopoulos, et al. 1999; Tenaillon, et al. 2016). By contrast, only one of the canonical hypermutator populations, Ara−3, shows an IS hypermutator phenotype. The rate of observed structural mutations in Ara−3 shows three different slopes. Ara−3 evolved an IS hypermutator phenotype very early in the LTEE. Around 30,000 generations, the IS rate intensifies, either due to genetic evolution, or as a consequence of stress induced by the citrate metabolic innovation that evolved around that time (Blount, et al. 2012; Blount, et al. 2020). Finally, the IS rate decreases around 45,000 generations. More than 100 mutations go fixation in the selective sweep at 45,000 generations in Ara−3, including mutations in the DNA repair genes *recR*, *recE, ligA, uvrA,* and *ybaZ*. The distinct IS rates observed in Ara−3 may, in part, reflect the relative frequency of deeply diverged, competing lineages in that population (Blount, et al. 2012), especially if those lineages had different IS transposition rates.

### Evidence of mutation biases based on gene orientation on the chromosome

It has been reported that genes on the lagging strand have higher mutation rates than genes on the leading strand in the DNA replication bubble, due to head-on collisions between the replication and transcription molecular machinery (Paul, et al. 2013). Based on such reports, we asked whether the LTEE metagenomics data showed evidence of gene-orientation mutation biases (Figure 4). Each distribution in Figure 4 is asymmetric over the replication origin. At the replication origin, one DNA strand switches from leading to lagging, while its complement switches from lagging to leading. This switch occurs because DNA replication is bidirectional, such that two replisomes move in opposite directions from the replication origin. Thus, the observed asymmetry over the replication origin is consistent with gene-orientation mutation biases. Indeed, the number of observed mutations significantly differs between genes oriented with or against the movement of the replisome, based on comparing the expected ratio of mutations to the observed ratio of mutations (binomial test: *p* < 10^−10^).

**Figure 4.**
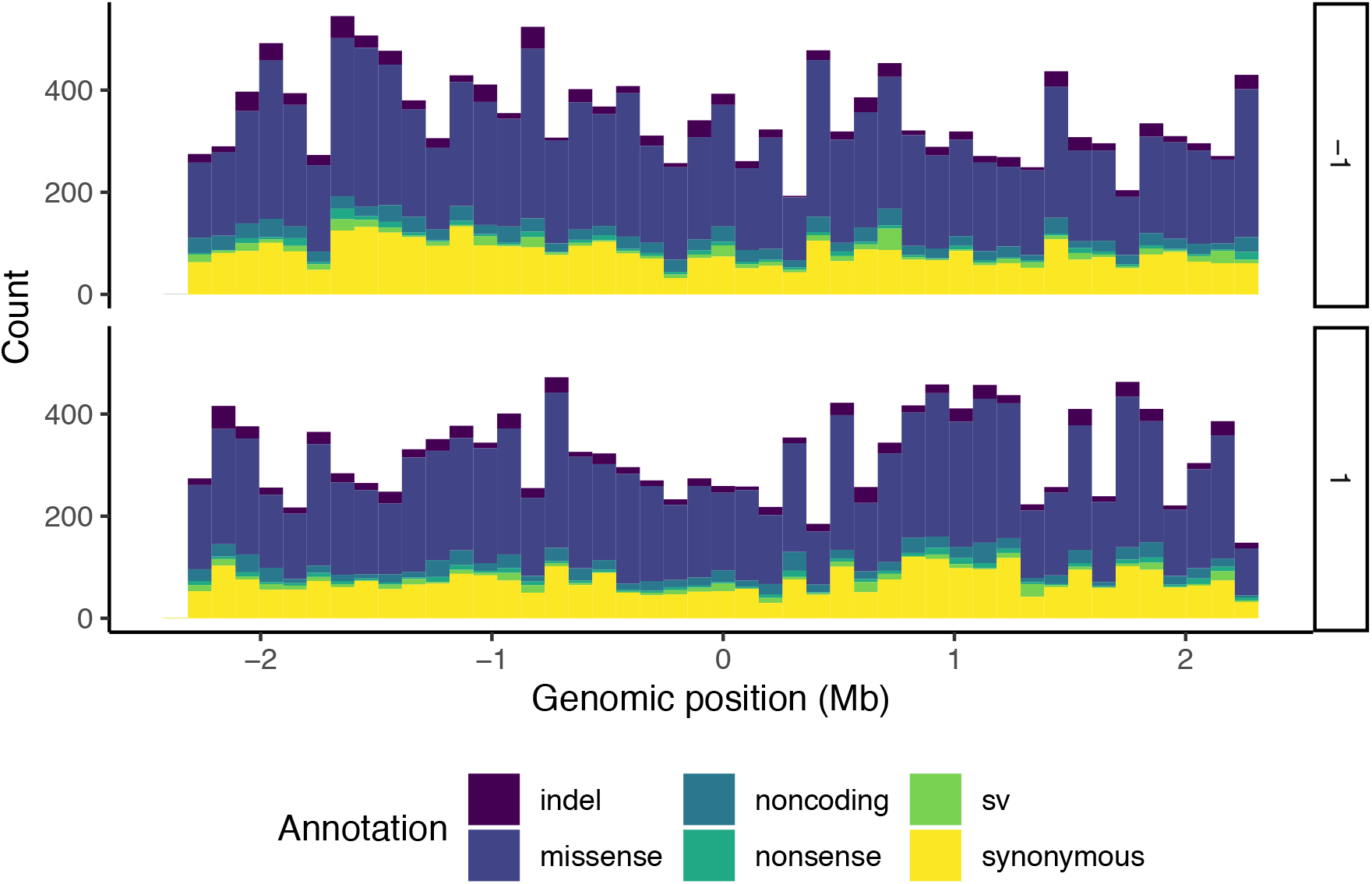
Evidence for gene-orientation mutation bias in the LTEE. Each panel shows the genomic distribution of mutations within genes, summed over all LTEE populations. The x-axis is the reference genome, centered on the replication origin, partitioned into 46 equally-sized bins of ~100 kb. The two panels show genes occurring on each of the two strands of the chromosome, with the arbitrary labels 1 and −1. Indels are in purple, missense mutations are in dark blue, noncoding mutations are blue green, nonsense mutations are sea green, structural variants (sv) are green, and synonymous mutations are yellow.

### The genomic distribution of observed mutations in Ara+3 shows a strong, symmetric wave pattern over the origin of replication

Multiple studies (Sharp, et al. 1989; Lang and Murray 2011; Foster, et al. 2013; Dillon, et al. 2018; Niccum, et al. 2019) have reported correlations between local mutation rates and distance from the origin of replication. One hypermutator LTEE population, called Ara+3, shows a symmetric wave pattern reflected over *oriC* (Figure 5). The wave in Ara+3 has a trough-to-peak ratio of ~25:75 (Figure 5). Excluding Ara+3, the genomic distribution of observed mutations summed over the remaining mismatch-repair deficit LTEE populations shows a weak wave pattern, while the populations with defects in *mutT* shows no evidence of the wave pattern (Figure 6).

**Figure 5.**
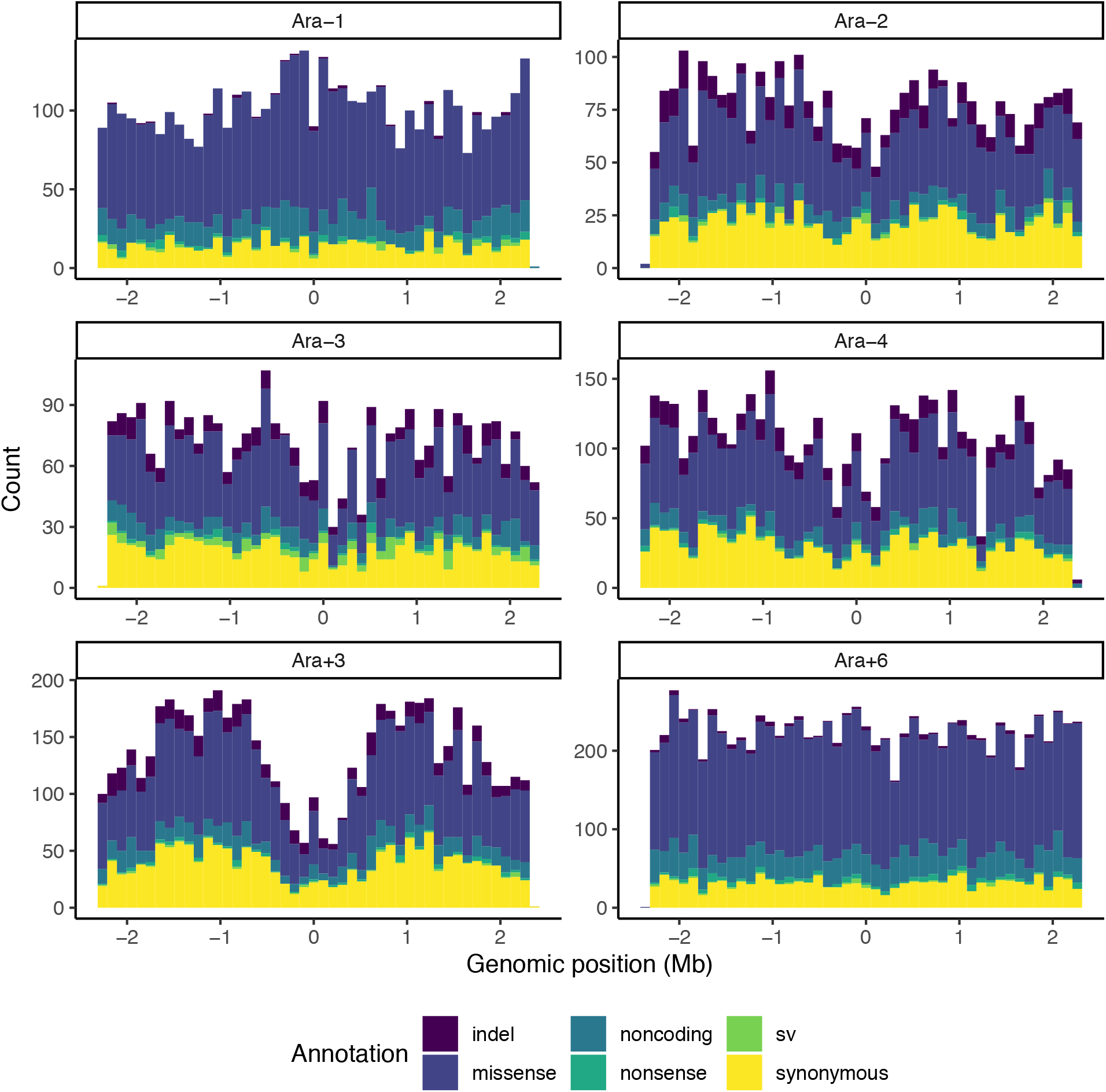
One hypermutator LTEE population, Ara+3, shows a strong wave pattern of mutation rate variation centered on the replication origin. Each panel shows the genomic distribution of mutations observed in each hypermutator LTEE population in the metagenomics data. The x-axis is the reference genome, centered on the replication origin, partitioned into 46 equally-sized bins of ~100 kb. Indels are in purple, missense mutations are in dark blue, noncoding mutations are blue green, nonsense mutations are sea green, structural variants (sv) are green, and synonymous mutations are yellow.

**Figure 6.**
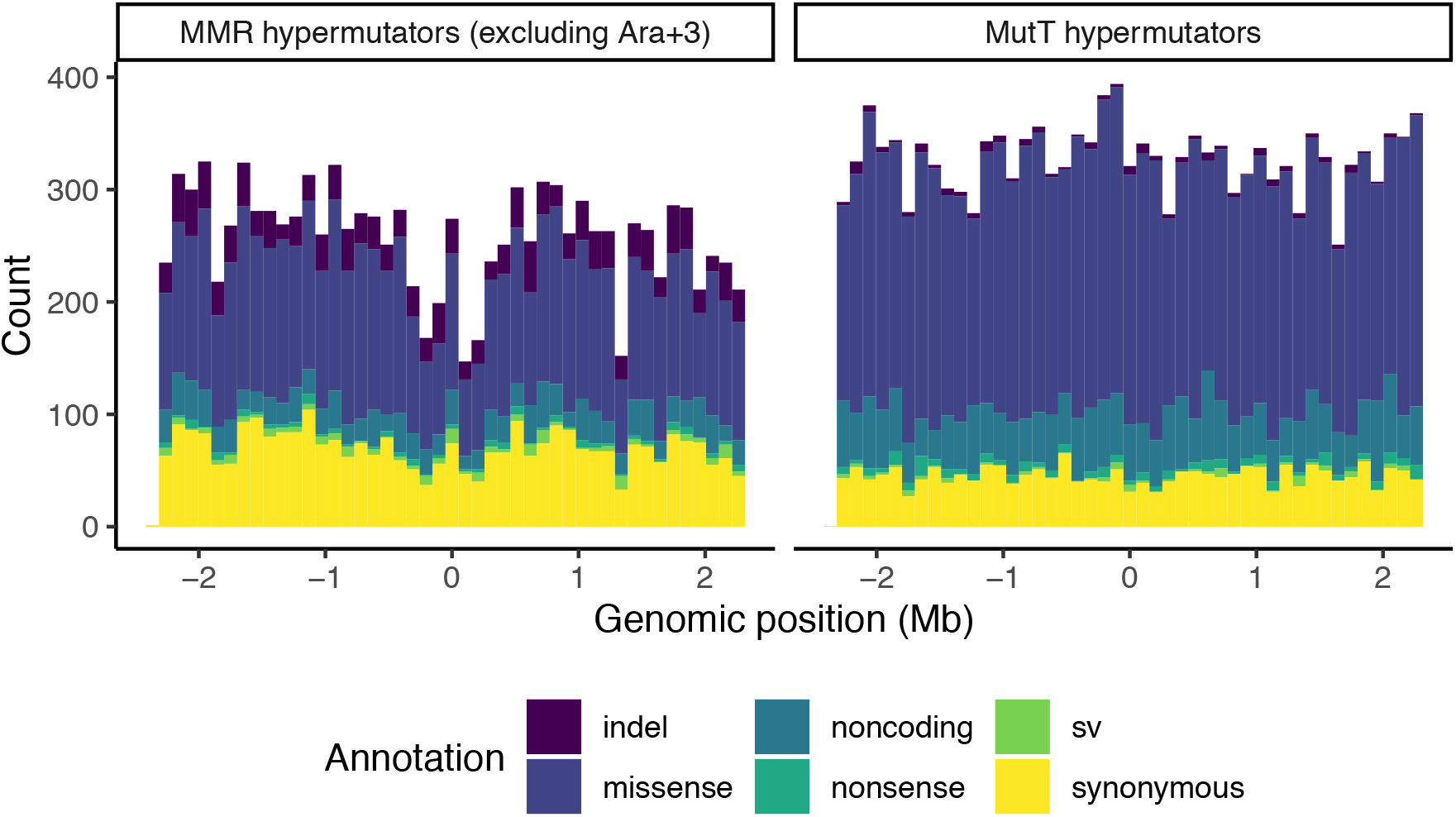
Mismatch-repair deficient LTEE populations (excluding Ara+3) show a weak wave pattern, while MutT-deficient LTEE populations show no wave pattern. The left panel shows the genomic distribution of mutations observed in Ara−2, Ara−3, and Ara−4. The right panel shows the genomic distribution of mutations observed in Ara−1 and Ara+6. The x-axis is the reference genome, centered on the replication origin, partitioned into 46 equally-sized bins of ~100 kb. Indels are in purple, missense mutations are in dark blue, noncoding mutations are blue green, nonsense mutations are sea green, structural variants (sv) are green, and synonymous mutations are yellow.

### Evidence for epistasis and historical contingency in the evolution of DNA topology

Why does a strong wave pattern only appear in Ara+3? Others have hypothesized that local chromatin structure affects local mutation rates (Foster, et al. 2013; Niccum, et al. 2019). Furthermore, DNA topology is an known target for selection in the LTEE (Crozat, et al. 2005; Crozat, et al. 2010). Therefore, we hypothesized that mutations in genes that affect DNA topology might affect the wave pattern. To test this hypothesis, we examined the timing and distribution of mutations in *topA, fis,* and *dusB* (*yhdG*), as alleles of these genes are known to affect DNA topology in the LTEE (Crozat, et al. 2005; Crozat, et al. 2010). We excluded synonymous mutations from this analysis.

All LTEE populations evolved missense, indel, or structural mutations in *topA, fis,* and *dusB* within the first 10,000 generations, except two: Ara+2, and Ara+3 (Figure 7). The timing and distribution of mutations in these genes across populations suggests epistasis and historical contingency (Good, et al. 2017). The early arrival times for mutations in these genes suggests that there is an early, limited window of opportunity for those mutations to go to fixation. Quantitative evidence comes from Ara+3, which has no missense, indel, or structural mutations in *topA, fis,* and *dusB* whatsoever, despite its strong hypermutator phenotype. The probability of this event is *p* = (1 − (*t* / *g*))^*n*^, where *t* is the effective mutational target size, *g* is the length of the chromosome (*g =* 4,629,812), and *n* is the number of observed missense, indel, and structural mutations in Ara+3 (*n* = 4,368). Given the wave pattern in Ara+3, the effective mutational target size of *topA*, *fis*, and *dusB* could be smaller than their combined physical target size (3,861 bp), say if they occurred in the trough of the wave. To take this into account, we partitioned the chromosome into bins, counted mutations per bin, and calculated the effective mutational target size by multiplying the physical target size (length) of *topA, fis,* and *dusB* by the number of mutations per base pair in their respective bins. These genes are significantly depleted of mutations in Ara+3, for bin sizes ranging from 100 kb to the entire chromosome (*p* < 0.05).

**Figure 7.**
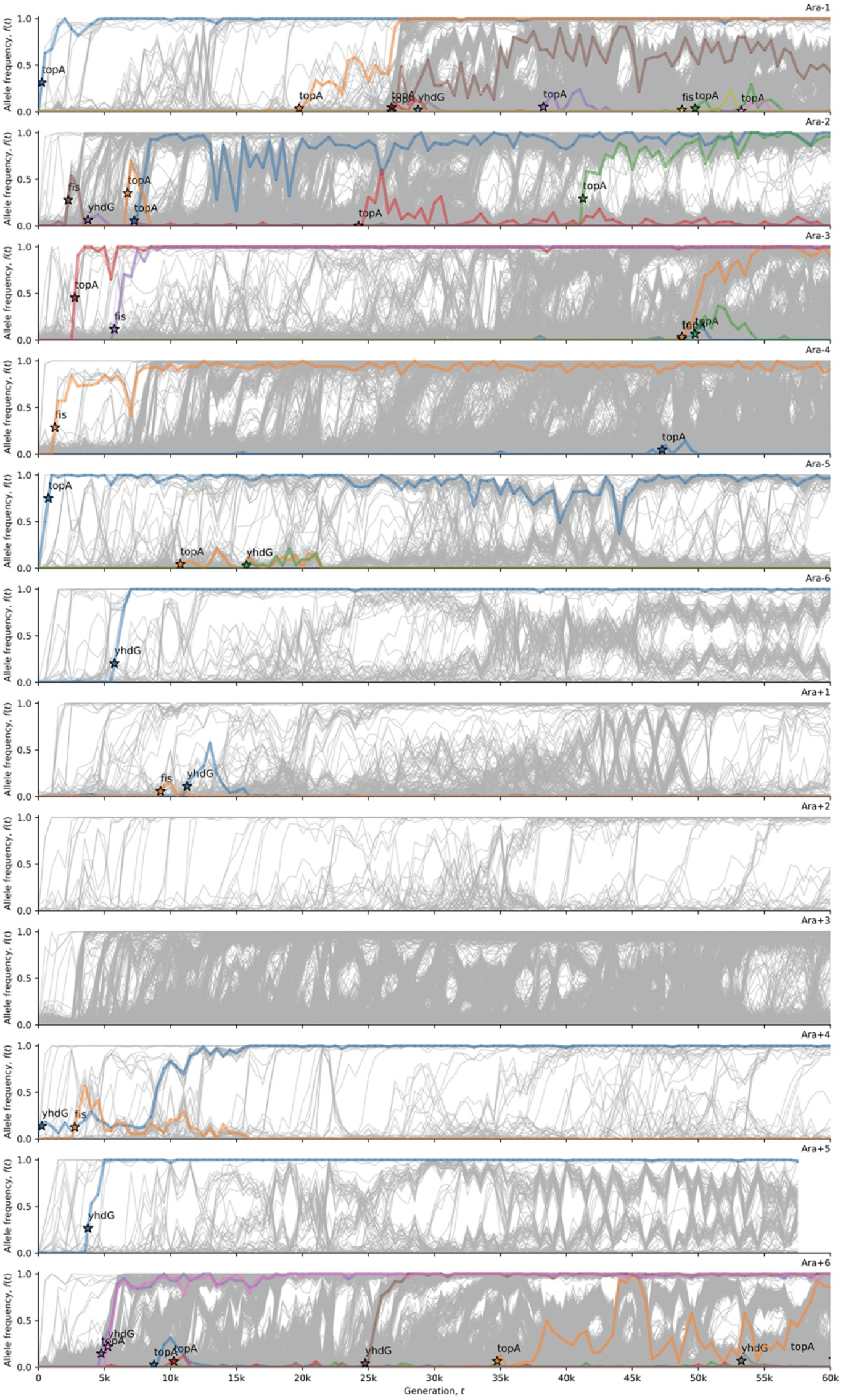
The strong wave pattern in Ara+3 anti-correlates with mutations (excluding synonymous changes) in the DNA topology genes *topA, fis,* and *dusB* (labeled as *yhdG*). This visualization uses computer code written by Good et al. (2017). The allele frequency trajectories for all observed mutations in the twelve LTEE populations are shown in grey. The allele frequency trajectories of *de novo* mutations in *topA, fis,* and *dusB* (excepting synonymous mutations) are colored and labeled in each population.

The distribution of synonymous mutations in *topA, fis,* and *dusB* across the LTEE populations is interesting (Supplementary Figure S4). A single, synonymous A312A substitution in *dusB* went to fixation at ~4000 generations in Ara+3, simultaneously with alleles in the mismatch repair genes *mutS* and *mutH* that apparently caused the early hypermutator phenotype in this population. No further synonymous mutations are observed in Ara+3. There is evidence of parallel evolution at this position in *dusB*. The same synonymous mutation in *dusB* occurs in Ara+6, and another synonymous mutation, one base pair downstream in the next codon, is the only synonymous mutation in *topA*, *fis*, or *dusB* observed in Ara−2 (Supplementary Figure S4). This parallelism suggests that positive selection may be acting on these synonymous variants. Overall, it is striking how few synonymous mutations in *topA, fis,* and *dusB* occur in the hypermutator LTEE populations, which implies that synonymous variants in these genes may not be evolving neutrally. Indeed, STIMS (Maddamsetti and Grant 2020) finds a significant signal of purifying selection on synonymous mutations in *topA*, *fis*, and *dusB* in Ara−1 and Ara−3 (one-tailed randomization test with 10,000 bootstraps; *p* < 0.0001).

### Synonymous nucleotide diversity in natural E. coli populations does not predict mutation rate variation in the LTEE

Finally, we used the LTEE metagenomic data to revisit previous work, which found that the distribution of synonymous mutations in the LTEE does not reflect patterns of synonymous variation among natural *E. coli* isolates (Maddamsetti, et al. 2015). During our reanalysis, we found that the usage of the Kolmogorov-Smirnov test in that paper was problematic. Therefore, we used Poisson regression to ask whether the estimates of synonymous nucleotide diversity θ_s_ published in (Martincorena, et al. 2012), when treated as gene-specific estimates of the point-mutation rate per base pair, predicts the distribution of synonymous mutations observed in the LTEE. A null model in which mutations occur uniformly over the chromosome (Akaike’s Information Criterion = 8529.6) fits the data far better than the θ_s_ model (Akaike’s Information Criterion = 9171.3). The same conclusion holds when we fit both models to Ara+3 (AIC = 2168.2 for null model versus AIC = 2190.8 for θ_s_ model).

## DISCUSSION

By examining the distribution of observed mutations over more than 60,000 generations of the LTEE (Good, et al. 2017), we find that mutation rates and biases have diverged idiosyncratically, despite identical abiotic conditions. One LTEE population, Ara+3, shows strong evidence of the wave pattern in mutation rate variation. Similar patterns have been seen in mutation accumulation experiments with MMR-deficient strains of *E. coli* as well as in *Vibrio* bacteria (Dillon, et al. 2018; Niccum, et al. 2019). Our result shows that genomic biases in mutation rates evolve dynamically on laboratory timescales.

The divergence in the rates, biases, and spectra of mutations across replicate populations in this simple long-term evolution experiment makes one wonder about the scope of natural variation in mutation rates, biases, and spectra. An evolution experiment with replicate mouse microbiomes has indicated that microbial evolution in the gut is probably characterized by long-term maintenance of intraspecies genetic diversity, including mutation rate polymorphism (Ramiro, et al. 2020). Phylogenomic studies have also found extensive evidence for horizontal gene transfer in DNA repair genes (Denamur, et al. 2000), which suggests that polymorphism in DNA repair genes may cause extensive natural variation in mutation and recombination rates within and across bacterial (meta-)populations and communities.

We find statistical evidence for historical contingency and epistasis in the evolution of DNA topology in the LTEE, and for Ara+3 in particular. These findings suggest a relationship between local DNA topology and local mutation rate variation, consistent with the experiments reported by Niccum et al (2019). These findings immediately suggest the need for experiments to test whether local DNA topology causes local mutation rate variation.

A comparison of synonymous genetic variation estimated from natural *E. coli* isolates to the distribution of observed synonymous mutations in the LTEE confirms the main result in earlier work (Maddamsetti, et al. 2015) using richer data, and is consistent with other reports as well (Lee, et al. 2012; Chen and Zhang 2013; Lynch, et al. 2016). In sum, gene-specific variation in synonymous nucleotide diversity θ_s_, estimated from natural isolates of *E. coli*, does not predict the genomic distribution of synonymous mutations observed in the LTEE. In any case, the other results that we have presented, in addition to prior reports (Foster, et al. 2013; Paul, et al. 2013; Jee, et al. 2016; Niccum, et al. 2019), strongly indicate that mutation rates vary over the *E. coli* chromosome.

These results add to the robust debate on the causes and consequences of mutation rate evolution. It is clear that a deeper understanding of the relationships among chromatin structure, genomic variation in mutation and recombination rates, and natural selection, and their consequences for short-term and long-term genome evolution, will be a fruitful goal for further research.

## MATERIALS AND METHODS

Pre-processed LTEE metagenomics data, and associated analysis and visualization code was downloaded from: https://github.com/benjaminhgood/LTEE-metagenomic. Analysis codes are available at: https://github.com/rohanmaddamsetti/LTEE-purifying-selection/blob/master/mutation-rate-analysis.R and https://github.com/rohanmaddamsetti/LTEE-purifying-selection/blob/master/metagenomics-library.R. We systematically examined DNA repair genes in *E. coli* (Eisen and Hanawalt 1999; Lee, et al. 2016; Deatherage, et al. 2018), as well as annotated DNA polymerases, and other proteins of the replisome. A table of these genes and their annotations are in Supplementary Data File 1. Datasets and analysis codes are available on the Dryad Digital Repository (DOI pending publication).

## ACKNOWLEDGEMENTS

We thank Richard Lenski for valuable discussions and advice. We thank Jeffrey Barrick for valuable discussions and comments on an earlier version of our manuscript. We thank Helen Murphy for valuable discussions, and thank Benjamin Good for making pre-processed LTEE metagenomics data and analysis scripts easily accessible for the research community. The LTEE that generated the bacteria we used in this study is supported by a grant from the National Science Foundation (currently DEB-1951307) to Richard Lenski and Jeffrey Barrick. N.A.G. was supported by the BEACON Center for the Study of Evolution in Action (NSF cooperative agreement DBI-0939454) and by Michigan State University.

## SUPPLEMENTARY INFORMATION

**Supplementary File 1: Annotated DNA repair and replication genes in** *Escherichia coli*.

**Supplementary Figure S1.**
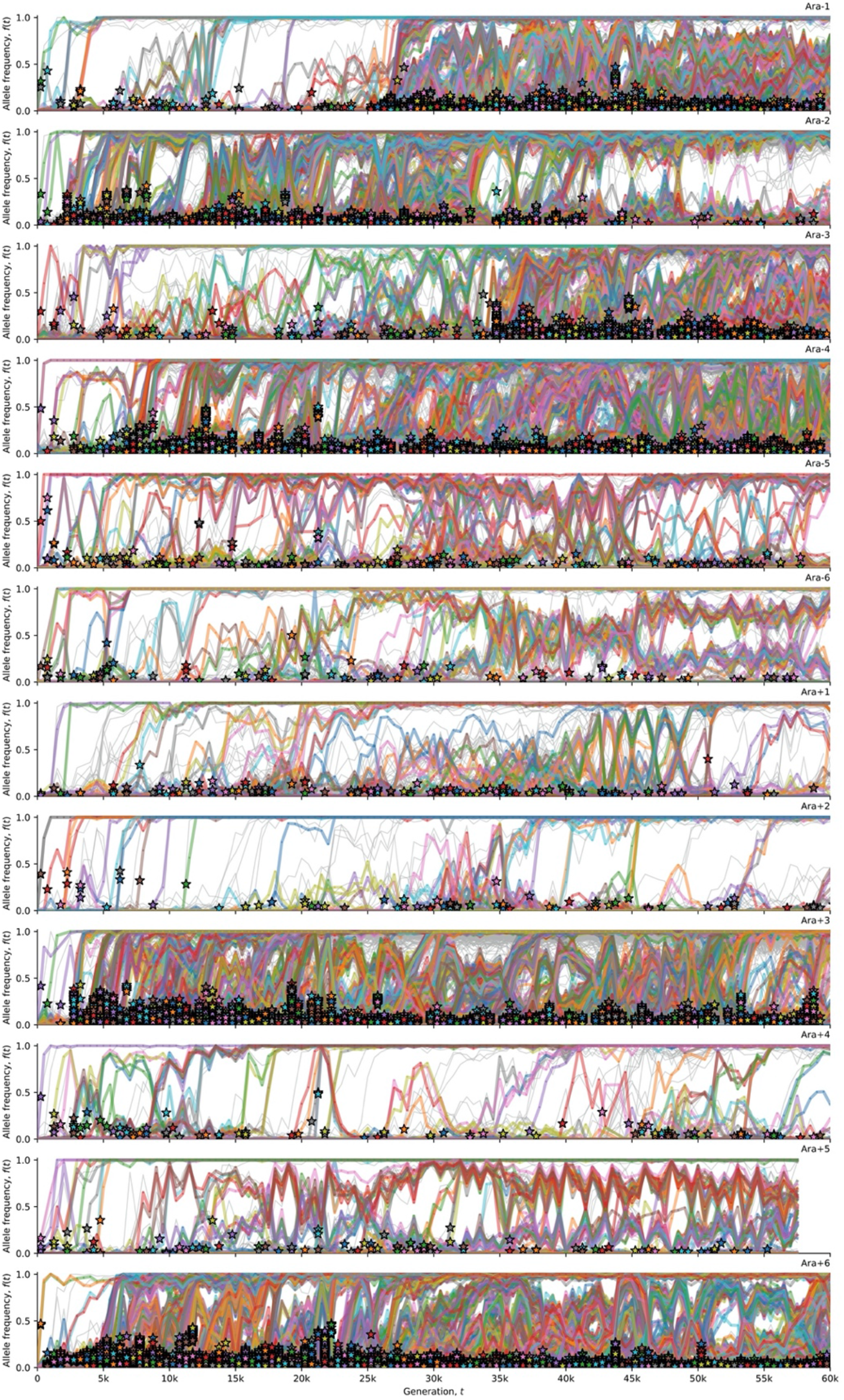
Allele frequency trajectories for all observed point mutations in the twelve LTEE populations. This visualization uses computer code written by Good et al. (2017). The allele frequency trajectories for all observed mutations in the twelve LTEE populations are shown in grey. Stars indicate the time (and allele frequency) at which mutations are reliably estimated to appear in the time series.

**Supplementary Figure S2.**
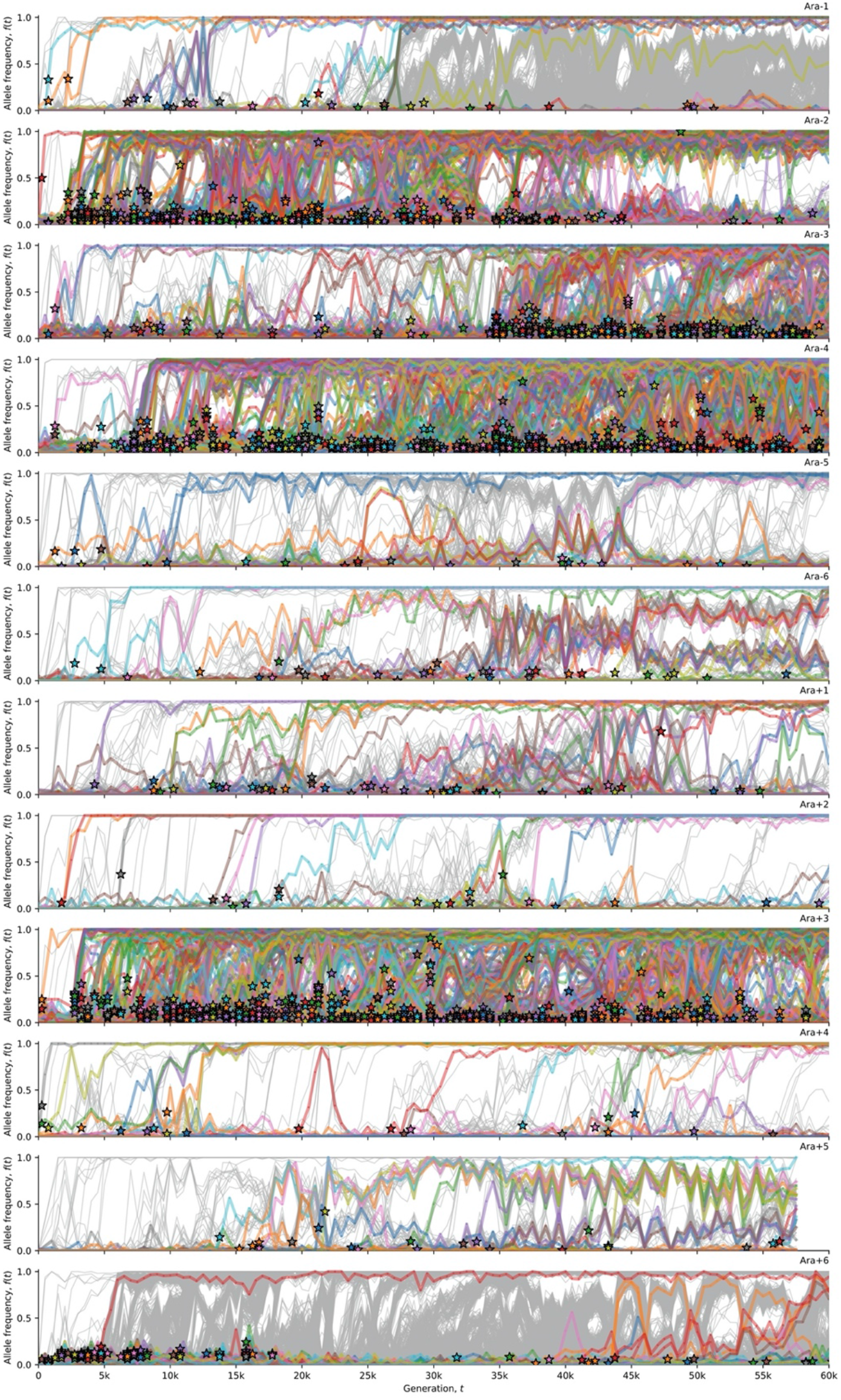
Allele frequency trajectories for all observed indel mutations in the twelve LTEE populations. This visualization uses computer code written by Good et al. (2017). The allele frequency trajectories for all observed mutations in the twelve LTEE populations are shown in grey. Stars indicate the time (and allele frequency) at which mutations are reliably estimated to appear in the time series.

**Supplementary Figure S3.**
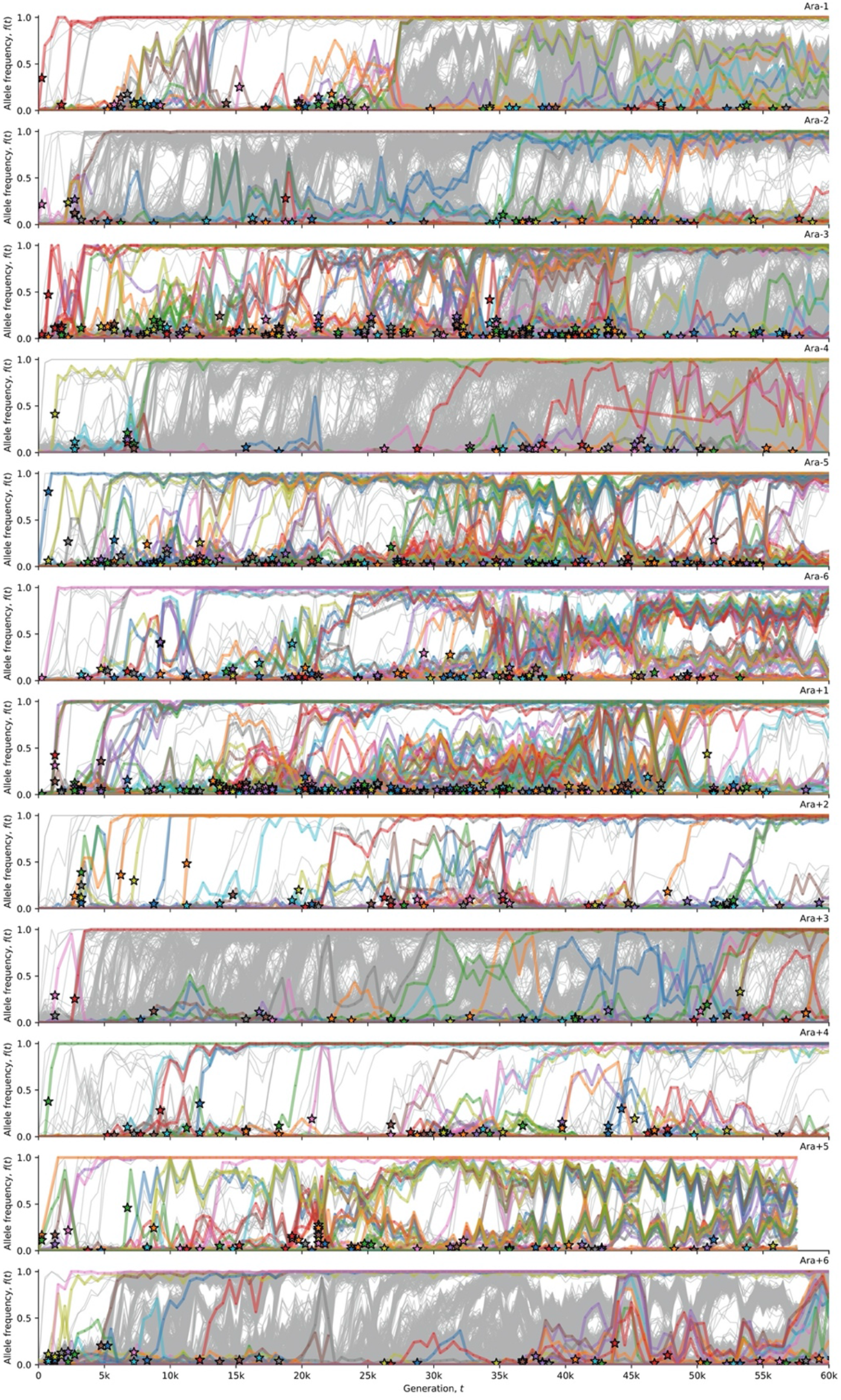
Allele frequency trajectories for all observed structural mutations in the twelve LTEE populations. This visualization uses computer code written by Good et al. (2017). The allele frequency trajectories for all observed mutations in the twelve LTEE populations are shown in grey. Stars indicate the time (and allele frequency) at which mutations are reliably estimated to appear in the time series.

**Supplementary Figure S4.**
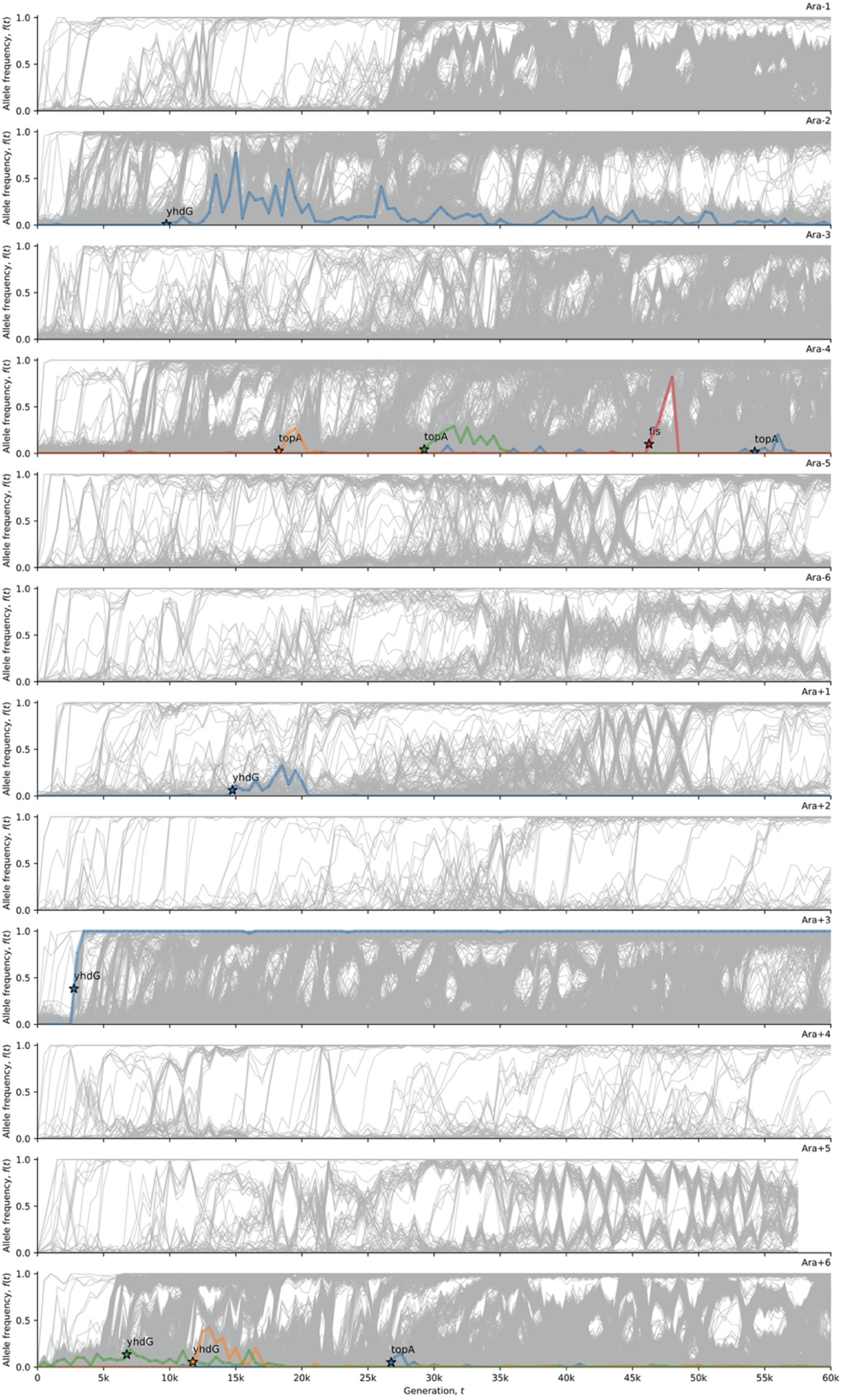
Synonymous mutations in the DNA topology genes *topA, fis,* and *dusB* (labeled as *yhdG*). This visualization uses computer code written by Good et al. (2017). The allele frequency trajectories for all observed mutations in the twelve LTEE populations are shown in grey. The allele frequency trajectories of *de novo* synonymous mutations in *topA, fis,* and *dusB* are colored and labeled in each population.

